# Machine Learning-based Identification and Characterization of High-risk Carriers of HTLV-1-Associated Myelopathy (HAM)

**DOI:** 10.1101/2025.03.17.643639

**Authors:** Md Ishtiak Rashid, Junya Sunagawa, Akari Matsuki, Asami Yamada, Toshiki Watanabe, Masako Iwanaga, Ki-Ryang Koh, Takafumi Shichijo, Masao Matsuoka, Jun-ichirou Yasunaga, Shinji Nakaoka

**Affiliations:** Graduate School of Life Science, Hokkaido University, Sapporo, Japan; Quantitative Life Sciences, The Abdus Salam International Centre for Theoretical Physics (ICTP), Italy; Faculty of Advanced Life Science, Hokkaido University, Sapporo, Japan; Department of Hematology, Rheumatology, and Infectious Diseases, Graduate School of Medical Sciences, Faculty of Life Sciences, Kumamoto University, Kumamoto, Japan; Kumamoto Rosai Hospital, 1670 Takehararamachi, Yatsushiro, Kumamoto, Japan; Department of Hematology & Oncology, St. Marianna University School of Medicine, Kanagawa, Japan; Department of Clinical Epidemiology, Nagasaki University Graduate School of Biomedical Sciences, Nagasaki, Japan; Nagasaki Genbaku Hospital, Nagasaki, Japan; Department of Hematology, Osaka General Hospital of West Japan Railway Company, Osaka, Japan

## Abstract

HTLV-1-associated myelopathy (HAM) develops in a part of HTLV-1-infected individuals while most of the individuals remain asymptomatic. This complicates the identification of HTLV-1 carriers at elevated risk. In this study, we integrated HTLV-1 proviral load and antibody titers against Tax, Env, Gag p15, p19, and p24 proteins in a machine learning (ML) framework to identify and characterize high-risk individuals likely to develop HAM. We stratified asymptomatic carrier samples employing an anomaly detection model. We further developed and validated classifier models capable of distinguishing three clinical subgroups, carrier, ATL, and HAM for assessing the anomaly carrier samples as unseen test data. With most anomaly carrier samples (∼76.47%) predicted as HAM, further statistical and interpretative analysis revealed the ‘HAM-like’ characteristics of the anomaly carrier samples indicating elevated risk. Additionally, significant heterogeneity in immune response was observed among other asymptomatic carriers. Our machine learning-based approach offers a novel and insightful tool for identifying and evaluating high-risk characteristics for HAM, providing a holistic view of the complex immune dynamics of asymptomatic carriers of HTLV-1.

## Introduction

Human T-cell leukemia virus type 1 (HTLV-1) is the first identified retrovirus known to cause chronic lifelong infections in humans [1]. Following infection, it causes adult T-cell leukemia-lymphoma (ATL), a form of blood cancer, and HTLV-1-associated myelopathy (HAM), a chronic inflammatory disease of the central nervous system. While most infected individuals remain asymptomatic, approximately 2 to 5% of HTLV-1 carriers develop ATL [2], and 0.25 to 4% develop HAM/TSP [3]. The annual national incidence rate of new HTLV-1 infections in Japan has been reported as 3.8 cases per 100,000 person-years [4]. The combined strategies of immune evasion and suppression allow HTLV-1 to remain in the host for a prolonged time without expressing symptoms which complicates the identification of carriers at higher risk of developing ATL or HAM [5][6][7]. ATL has been reported to occur in adults at least 20 to 30 years after the infection [8] and for HAM, the latent period ranged from 4 months to 30 years [9]. While it remains unknown why some HTLV-1 carriers develop the disease, it is of interest to develop methods that can identify high-risk asymptomatic individuals.

For HTLV-1-infected individuals, the diversity of the infected cells and immune responses is influenced by the interaction between the virus and the host’s immune system. A disruption of the equilibrium between viral persistence and host immunity can trigger the onset of associated diseases. Distinct immunological responses may arise from impaired immune regulation against the HTLV-1 infection [10]. As reported previously, HTLV-1 infection evokes both cellular and humoral immunity [11][12][13]. Therefore, immunological markers can be instrumental in evaluating the risk of associated diseases. Although proviral load (PVL) has been considered a risk factor, its interpretation in asymptomatic carriers is still challenging due to individual variations and lack of definitive thresholds for disease risk [14][15]. Additionally, some studies reported the PVL quantity remains constant over several years, regardless of clinical manifestation [15][16][17][18]. However, antibody profiling, when combined with other markers like PVL, has been reported as useful for distinguishing asymptomatic carriers from ATL or HAM patients [19][20][21][22] and potential for predicting disease progression. Moreover, serum-based testing is simpler and cheaper than genetic analysis. Hence, analyzing antibody responses can provide a quantitative and specific method for identifying the infected individuals at elevated risk.

In our previous study using a modified Luciferase immunoprecipitation system (LIPS) assay, we screened patients for antibodies against each of the viral proteins, HTLV-1 Gag proteins (p15, p19, p24), Env, and Tax. Along with multivariate analysis of the antibody titers followed by targeted sequencing, we identified carriers at high risk for ATL. We also listed important factors for separating the HAM subgroup [21]; however, this finding might overlook the possibility that some carriers are already showing similar antibody responses to HAM/TSP and thus need to be explored in more depth. Although longitudinal studies are invaluable for observing the disease progression over time, it is still challenging given a prolonged latency period. Moreover, considering the treatment effects, it may not fully capture the hidden patterns within the data. Cohort studies usually maintain high data quality and standardized data collection protocols. This reduces noise and inconsistencies, making ML models more compatible and generalizable [23]. Machine learning (ML) models surpass the traditional multivariate statistical methods by handling complex, high-dimensional data and uncovering non-linear patterns. Based on clinical data alone ML models offer a robust alternative tool within data-rich contexts [24][25][26][27].

In the present study, we developed a two-tiered ML-based framework integrating antibody titers to HTLV-1 PVL and Tax, Env, along with the immunogenic mature Gag p15, p19, and p24 proteins to identify and characterize the asymptomatic carriers with a higher likelihood of developing HAM. Collectively our results show that the ML-based approach can be effective in risk prediction and early intervention of HAM that might otherwise remain undetected through conventional diagnostic approaches.

## Results

### Detection of Potentially High-Risk Carriers

At the data preprocessing step, before applying ML models to our dataset, we handled the multicollinearity issue [28] by excluding Gag p19, based on the highest VIF score (see **Determination of Key Variables** in **Method** and Supplementary Table S1A and B for details). Out of 264 asymptomatic carriers, the Isolation forest model detected 17 carrier samples as anomalies, which we labeled as anomaly carriers (AC). One carrier who later developed HAM/TSP (CDH) was also identified as an anomaly and included in the AC group, suggesting that the Isolation forest model effectively filters carrier samples with a possible risk of HTLV-1-associated diseases. For details about the algorithm of Isolation forest and choice of hyperparameters, please see **Anomaly Detection by Isolation Forest Algorithm** in **Methods**.

Next, we compared the performance of four ML models based on PRAUC scores to classify samples into 3 subgroups: non-anomaly carrier, HAM/TSP, and ATL. We chose the Random forest model (RF) for its superior performance (See **Classification Modeling** in **Method** for details) [Supplementary Table S3]. The RF classifier model was used to predict the holdout set of AC samples (n=17). Interestingly, the classifier predicted the majority of AC samples (n=13 out of 17) as HAM/TSP [Figure 1]. The one CDH sample was also predicted as HAM/TSP by the classifier model. Moreover, in the case of 17 anomaly carrier samples, all the feature values showed a stronger positive correlation with their predicted probabilities of HAM/TSP except PVL [Supplementary Table S4].

**Figure 1:**
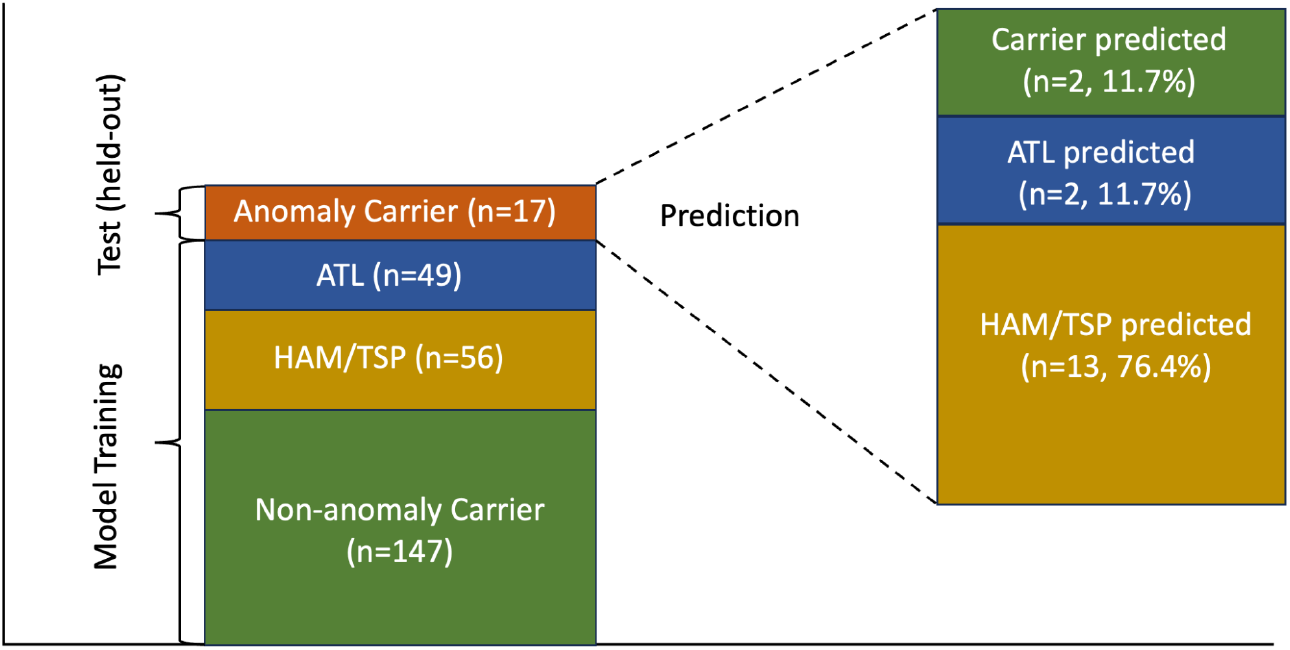
Illustration of the prediction results of anomaly carrier samples by the random forest classifier model. The left bar shows the training and test data for the classifier model. The model was trained on three sample groups and predicted the anomaly carrier samples as unseen test data. The anomaly carrier samples (n=17) were classified into three prediction groups: Around 76.4% of the anomaly carrier data were predicted as HAM/TSP, whereas only 11.7% and 11.7% of the samples were predicted as ATL and carrier respectively (shown on the right bar).

Results of Partial Least Squares (PLS) distribution of sample groups further revealed that AC samples were localized near the HAM/TSP cluster [Figure 2]. This result combined with the classification result where most AC samples predicted as HAM/TSP, suggests that AC samples share significant similarities with HAM/TSP samples, further supporting our hypothesis that AC samples exhibit ‘HAM-like’ characteristics.

**Figure 2:**
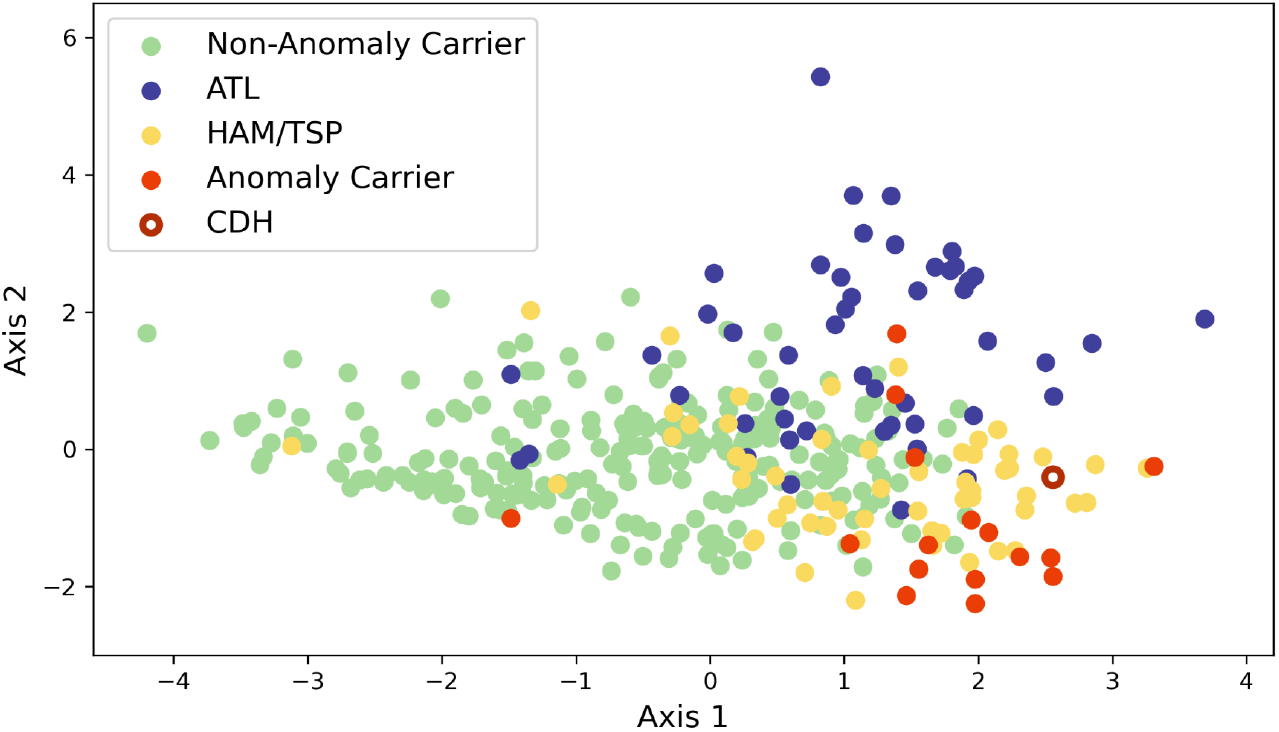
The PLS plane shows the distribution of all 369 samples from non-anomaly carriers (green), ATL patients (blue), HAM/TSP patients (yellow), anomaly carriers (red), and CDH (dark red as edge color and white-centered). Each dot on the plot represents an individual sample. This plot depicts the clustering of the sample groups based on the analyzed variables, where anomaly carrier samples were positioned near the HAM/TSP cluster.

### Comparison of Feature Value Distributions among the Subgroups

Figure 3 compares the feature values of 4 subgroups. Between AC and non-anomaly carrier comparison, all features in the anomaly carrier were significantly higher. On the contrary, we found no statistically significant difference between AC and HAM/TSP for PVL (p = 1.0), Tax (p = 1.0), Gag p15 (p = 0.34), Gag p19 (p=0.65) and Gag p24 (p = 0.89) except Env (p=0.0048) (for details about the statistical tests, please see **Boxplot Visualization and Statistical Tests** in **Methods)**. These observations collectively indicate that AC samples display a high degree of similarity with HAM/TSP. Consistent with previous research [10], we found antibody responses to the immunodominant proteins (Env, Tax, Gags) higher in HAM/TSP patients (n=56). Conversely, PVL in ATL patients was significantly higher than in all other subgroups [Figure 3, Supplementary Table S6], which is consistent with previous studies [4][29][30].

**Figure 3:**
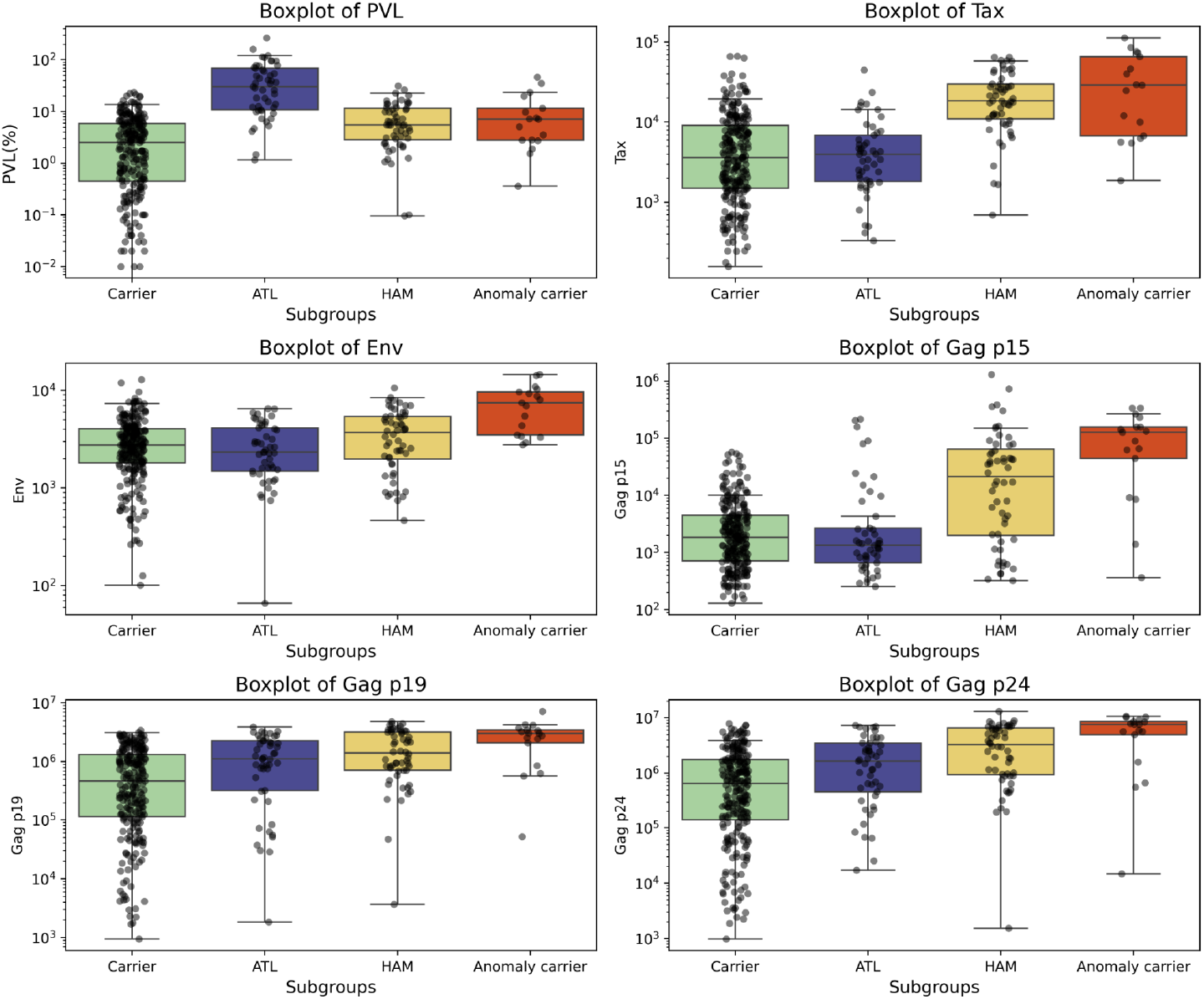
The boxplots collectively illustrate the distribution of PVL and Antibody titers to HTLV-1 antigens Tax, Env, Gag p15, Gag p19, and Gag p24 across different clinical subgroups: non-anomaly carrier (green), ATL (blue), HAM/TSP (yellow), and anomaly carrier (red). The individual data points overlaid on the boxplots show the actual distribution and density of the data.

### Exploring Biomarkers for Anomaly Carrier Detection

To determine the crucial factors that characterize the anomaly carrier and HAM/TSP, SHapley Additive exPlanations (SHAP) analysis was performed. SHAP analysis is an interpretable machine learning framework that can assess the impact of each feature on the classification of each class (non-anomaly carriers, ATL, HAM/TSP, and anomaly carriers). Figure 4 shows the SHAP bar plot from the ETC classifier which performed the best [Supplementary Figure S6]. We found Tax is the most important feature for HAM/TSP, and Gag p24 for anomaly carriers, followed by Gag p15 and Env. Although the prediction above indicated high similarity between anomaly carriers and HAM/TSP subgroups, the ranking of feature importance differs qualitatively. Gag p15 and Env are influential features in anomaly carriers but their relative rankings are not uniformly elevated in HAM/TSP. Also, Tax exhibits a lower ranking in the anomaly carrier.

**Figure 4:**
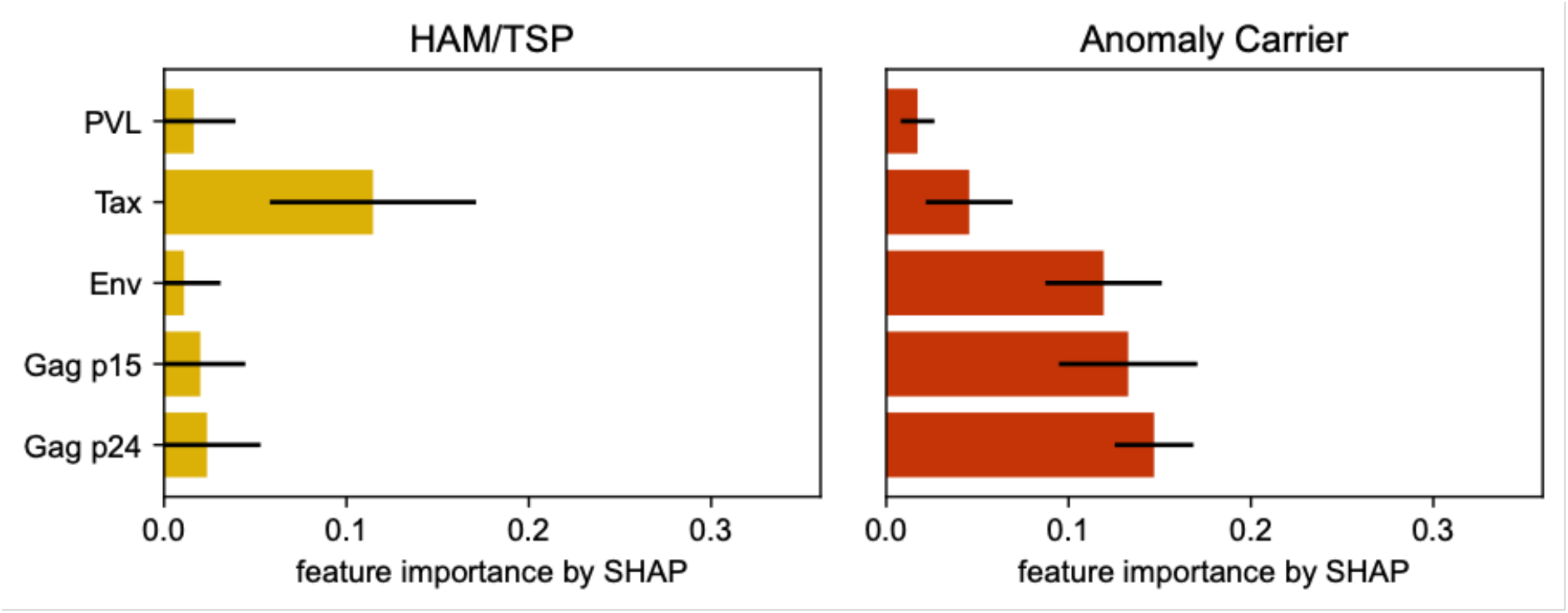
SHAP feature analysis of HAM/TSP and anomaly carriers. Each bar shows an absolute median value of 1000 iterations for different random seeds and the black line represents standard deviation. SHAP analysis for non-anomaly carriers and ATL is in Supplementary Figure S7.

## Discussion

In this study, we developed a machine learning-based approach to capture HTLV-1 carriers at elevated risk of HAM progression. The Isolation forest anomaly detection algorithm identified a subgroup of anomaly samples from the asymptomatic HTLV-1 carrier population. Further characterization through classifier prediction and statistical analysis revealed that the anomaly carrier samples closely resemble the characteristics of HAM, suggesting a similar disease trajectory. Additionally, different patterns of antibody response were observed among the asymptomatic carriers and other clinical subgroups which enabled us to further investigate the risk factors. Finally, we utilized SHAP for comparative feature analysis among the sample groups (non-anomaly carrier, anomaly carrier, ATL, and HAM) to identify the key driving features that characterize each subgroup and contribute to the disease progression.

The main aim of this study was to shed light on asymptomatic carriers who are at a high risk of progressing HAM onset. With most of the anomaly carrier samples being predicted as HAM by the RF classifier [Figure 1], our hypothesis was further supported when the purposely included CDH sample in the carrier population was also identified as an anomaly and subsequently predicted as HAM. The potential similarities in the underlying profiles of the anomaly carrier samples are also reflected in their clustering near the HAM samples [Figure 2]. All features were significantly higher in anomaly carriers compared to non-anomaly carriers [Figure 3]. Elevated antibody responses in anomaly carriers might reflect the immune response have higher activity during disease progression. Interestingly, we found that only anti-Env antibody titer in anomaly carriers differed significantly from those of HAM [Figure 3, Supplementary Table S6], whereas other features showed no significant differences. Env is one of the structural proteins of a virion and is necessary for cell-to-cell transmission. Thus it is a primary target of the antibody response [31, 32, 33, 34]. Furthermore, elevated anti-Env antibody responses have been associated with HAM patients in several studies, which supports our result [10, 19, 35, 36]. A novel implication is that, before onset, the rate of progression accelerates, as evidenced by the increased antibody levels. In HAM, the immune response is fully engaged; however, in progressive asymptomatic carriers, this saturation has yet to be achieved [19]. This phase might represent a snapshot of dynamic host-virus interaction where these rising antibody titers likely reflect the heightened viral activity and the immune system’s escalating response as the disease advances toward clinical manifestation. Ultimately, a saturation point is reached at the onset of the disease, where antibody levels level off as the immune response shifts into a steady-state phase. This might be well reflected in feature analysis, where the SHAP value of Env is relatively high in the anomaly carrier, but not in HAM and non-anomaly carrier [Figure 4, Supplementary Figure S7].

We found Tax to be the predominant feature of HAM, consistent with findings from multiple studies. [19, 37]. Furthermore, prior studies have reported significantly higher antibody responses to Env and Gag proteins in HAM patients reinforcing their potential role in HAM patients [10, 19]. It is known that during infection, Gag and Env proteins are initially unpolarized in isolated T cells and accumulate at the cell-cell junction upon contact. Gag protein is subsequently transferred from HTLV-1-infected T cells to uninfected T cells [38]. Aligning with these previous observations, we interestingly found the feature values of Gag p15, p24, and Env of anomaly carrier samples exhibited a significant inverse relationship with their anomaly scores, i.e., higher feature values correspond to higher anomaly levels [Supplementary Table S2, Supplementary Figure S5][39]. Assessment of humoral immunity to Gag demonstrates potential as a biomarker for detecting high-risk individuals. In our study, we succeeded in suggesting that Gag p15 protein has some important function that may lead to developing HAM onset [Figure 3 and 4], however, we avoid attributing our result to some implications about Gag p15; further research is required to identify the specific function of these mature Gag proteins (p15, p19, and p24). It is noteworthy that, although the SHAP value of both Gag p15 and p24 falls within the high-ranking features that characterize anomaly carriers, we opted to exclude the interpretation of Gags (p15, p24) due to their inconsistent contribution patterns observed across the multiple classifiers employed in this study [Figure 4 and Supplementary Figure S7].

Identifying the risk for developing HAM onset is challenging compared to other HTLV-1-associated diseases. In the case of ATL, for example, the risk can often be characterized by the changes in the clonality of infected cells, since a single clonal infected cell expands during the viral progression. Also, several driver mutations are reported to stimulate malignancy, thus leading to the survival of pathogenic cells and outcompete other infected cells towards monoclonal proliferation [40]. While these promising markers can detect risks of ATL onset, HAM is less described for early diagnosis, due to the nature of its slow progression [41]. Moreover, complicated host immune responses against infected cells vary widely between patients with different lifestyles, which makes the prediction more difficult [42]. Having anti-Env at the top of the list, elevated antibody titer might be a key observation for evaluating disease progression.

Of interest is the significant heterogeneity in immune response among the asymptomatic carriers in our study. Surprisingly, antibody responses (against Env, Tax, Gags, and PVL) in many asymptomatic carriers were observed at the same elevated level as that of HTLV 1-related diseases (ATL and HAM). This finding led us to our initial hypothesis to detect high-risk asymptomatic carriers (i.e., anomaly carriers) who are likely to progress to disease onset. Although heterogeneity seems to be obvious when considering the various lifestyle backgrounds of patients, it is noteworthy to confirm it based on our large number of asymptomatic carrier data.

Our work acknowledges some limitations. First, we don’t have information on anomaly carriers whether they develop HTLV-1-related diseases or not in the future except for one sample who was diagnosed as HAM later on (CDH). To fully evaluate the prediction and the hypothesis of our result especially for HAM/TSP, further data accumulation would be critical (a prospective study like [14]). Second, little is known about the relationship between the antibody titers and the host immune defense as mentioned above. For the data from the LIPS assay to be used as a clinical diagnosis, these interplays should be explored in more depth. Furthermore, inconsistent results in antibody titers from previous studies have discouraged clinical application, which makes it difficult to choose consensus cutoff values for disease distinction [41]. While cross-validation offers robust internal validation, we acknowledge the necessity of an independent dataset for evaluating the model’s generalizability. Future research should aim to include external validation cohorts to confirm our model’s applicability.

## Methods

### Ethics statement

This study was performed in accordance with the Declaration of Helsinki and was approved by the Ethics Committees of Kumamoto University (accession numbers: G489, G499, and E2214). Written informed consent was waived because of the retrospective design. Consent for publication was obtained from all patients.

### Study Population

The data used in this study was published previously by Yamada et al. [21]. PVL and antibody titer data (non-time series) were collected against HTLV-1 antigens Tax, Env, Gag p15, p19, and p24 using LIPS assay. Our previous study reported 24 carriers who later developed ATL (CDA) were considered as ATL in this study. We also had only one carrier who later developed HAM/TSP (CDH) and it was purposefully included in the carrier population. Therefore, we focused our study on 264 asymptomatic carriers, 49 ATL, and 56 HAM/TSP patients.

### Determination of Key Variables

Initially, Spearman’s rank correlation revealed a significant correlation between Gag p19 and p24 [Supplementary Figure S2]. To address the multicollinearity issue and choose the variables to use in the ML analysis, the Variance Inflation Factor (VIF) score was used [28]. See Supplementary Tables S1A and S1B.

### Anomaly Detection by Isolation Forest Algorithm

For identifying potential outliers or anomalous data points from the asymptomatic carrier population (n=264), we applied the Isolation Forest algorithm, an unsupervised machine learning technique based on decision trees. For each datapoint (sample), the following process is repeated until the datapoint is isolated:

1. Randomly select a feature (e.g. PVL)
2. Randomly choose a threshold between the maximum and minimum values of the selected feature (e.g. PVL=0.1) and divide the data points below and above the threshold.

The key idea is that data points with anomalous feature values are likely to be isolated with only a few iterations. The algorithm constructs an ensemble of isolation trees for a given dataset and uses the path length from the root to the leaf to determine the anomaly score. Given *m* is the number of data points, the anomaly scores *s* for a datapoint *x* is defined as

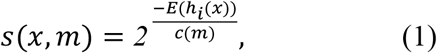

where *h*_*i*_(*x*) represents the path length for the *i* -th isolation tree, *E*(*h*_*i*_(*x*)) = ∑_*i*_ *h*_*i*_(*x*) denotes the average path length across the ensemble of isolation trees,

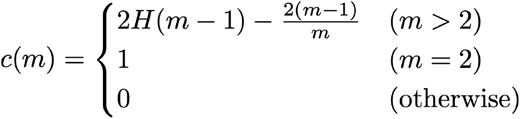

is the average path length for a dataset with m points, utilized as a normalization factor [43], and *H*(*k*) is the harmonic number. The sklearn implementation of the decision function of Isolation Forest yields negative anomaly scores, where lower (negative) scores indicate potential anomalies [39].

By applying a cutoff threshold at -0.05 to the anomaly scores of the Isolation forest, we isolated the anomaly data points for further investigation [44]. This threshold was strategically chosen to capture approximately 5% of the most extreme anomalies (inversely corresponding to the 95^th^ percentile of the normal data distribution) from our carrier population [Supplementary Figure S4]. Since around 4% of the carriers develop HAM/TSP [3, 45], we aimed to mirror this proportion.

The resulting anomaly carrier samples were then removed from the carrier data and considered as a holdout test set (unseen data) for further classification analysis. The remaining non-anomaly carrier, ATL, and HAM/TSP samples were used for training and cross-validation of the classifier models. Additionally, the feature values of the anomaly carrier samples were tested for Spearman correlation analysis with their anomaly scores. The difference between the sample groups was evaluated by plotting all the samples in a PLS plane.

### Classification Modeling

During data preprocessing, we randomly downsampled 100 carriers since the dataset was highly imbalanced. In this study, we employed the One-vs-Rest (OvR) approach to address the multiclass classification problem. This approach breaks down the multiclass classification into multiple binary classification tasks, where one classifier is trained for each class against all others. To determine the best-performing model, we evaluated four different classifiers: Random Forest classifier (RF), XGboost Classifier (XGB), Extra Trees Classifier models (ETC), and Support Vector Machine (SVM). Nested cross-validation (CV) was used to ensure robust performance evaluation and avoid overfitting. Particularly, an outer cross-validation loop was used to assess the model performances, while an inner loop was used to optimize the hyperparameters of each classifier using GridSearch.

In the inner loop, within each training set of the outer cross-validation, a 3-fold cross-validation was conducted for hyperparameter tuning and the combination that maximized the classifier performance based on the overall mean Area Under the Precision-Recall Curve (PRAUC) score was selected [46]. This ensured optimal parameter selection for each model. The model’s generalization performance was evaluated in the outer loop using a 5-fold cross-validation. For each fold, the dataset was partitioned into training and validation sets. The optimal hyperparameters identified in the inner loop were applied to train the classifier on the training subset, and the model’s performance was subsequently tested on the reserved test set. This process was repeated across all folds, and the average performance metrics were reported. Finally, the best classifier identified through nested CV was subsequently trained on the entire training dataset from the outer cross-validation and used to predict outcomes on the holdout test set (anomaly carrier samples). The predicted probability of HAM/TSP among the anomaly carrier samples (holdout test set) was calculated, followed by a correlation analysis of the predicted probabilities and their feature values. The workflow is depicted in [Supplementary Figure S1]. For the classification models performed in this study, the implementation available in the sklearn library was used [39]

### Boxplot Visualization and Statistical Analysis

We employed a combination of visual and statistical methods facilitating an initial comparison of the feature distributions among different sample groups including anomaly carriers. The Kruskal-Wallis test was performed, with a significance level set at α = 0.05. P-values were adjusted for multiple comparisons using the Bonferroni correction method for Dunn’s post-hoc analysis to maintain the overall type 1 error rate. The statistical analysis was performed using the Python Scipy package [47, 48].

### Interpretation with SHapley Additive exPlanations (SHAP) analysis

As an approach to interpreting the model’s behavior, the Shapley Additive exPlanations (SHAP) framework was used [49, 50]. It provides the SHAP value for each feature for all samples and explains how much an increase in each feature value can affect the predicted probability for each clinical subgroup (non-anomaly carriers, ATL, HAM/TSP, and anomaly carriers). A higher SHAP value indicates a greater impact on the classification of a sample into a specific subgroup, while a lower SHAP value corresponds to a smaller impact. In this section, four classifiers (RF, ETC, XGB, and SVM) were explored for their performance in terms of PRAUC using nested cross-validation and were calculated for 1000 different random seeds (i.e., different values for parameter random_state). Different random seeds are considered in this study because we wanted to extract the SHAP value which is consistent whenever the randomized manipulation during the learning process is different. This allows us to evaluate the results with a high degree of confidence. For each random seed, hyperparameters were optimized on all data without cross-validation by GridSearch and used for calculating SHAP value. The absolute median of the SHAP value from all samples was collected for 1000 random seeds, and then the absolute median value and its standard deviation were calculated for visualization. Specifically, KernelSHAP was applied for all classifiers in a SHAP python package (Version 0.45.1) [50].

## Data Availability

The dataset used for this study is not publicly available due to patient privacy concerns but may be made available from the corresponding author on request. All codes and files associated with this study are deposited and publicly available in the GitHub repository: https://github.com/petadimensionlab/HTLV1-machine-learning

## Acknowledgment

This work was supported by Japan Agency for Medical Research and Development, Grant/Award Number 21fk0108088h0003 and 22gm1710004h0001, Japan Science and Technology Agency (JST) Grant Number JPMJCR23J4, and JST Moonshot R&D Grant Number JPMJMS2021, JPMJMS2024-9 (to S.N.).

## Supplementary Information

### Supplementary Tables

**Supplementary Table S1A:**
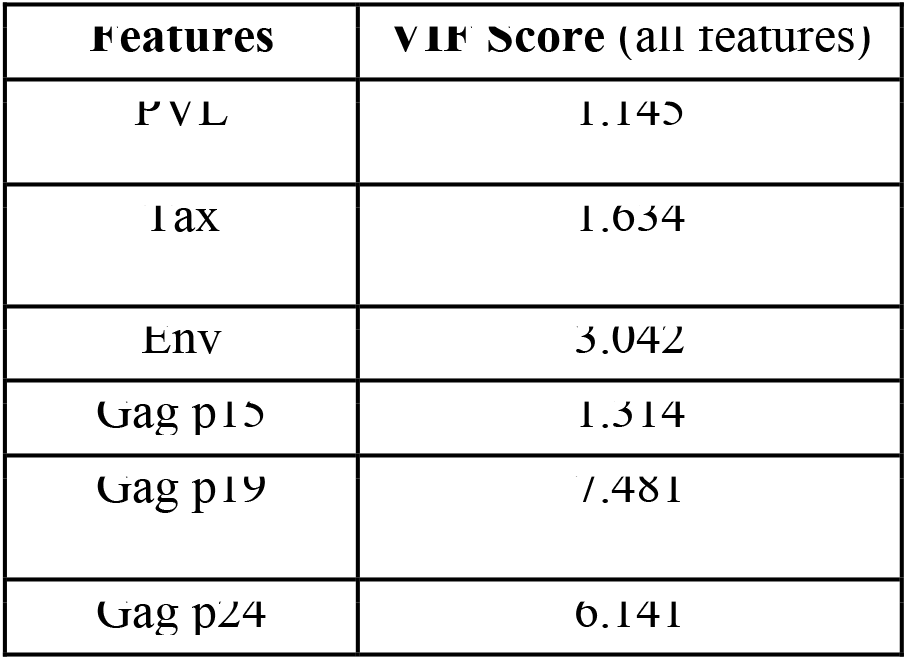
This table represents the feature values with the respective VIF scores. Gag p19 showed the highest VIF score of 7.480 for Gag p19, indicative of substantial multicollinearity. To mitigate this, Gag p19 was removed from the dataset.

**Supplementary Table S1B:**
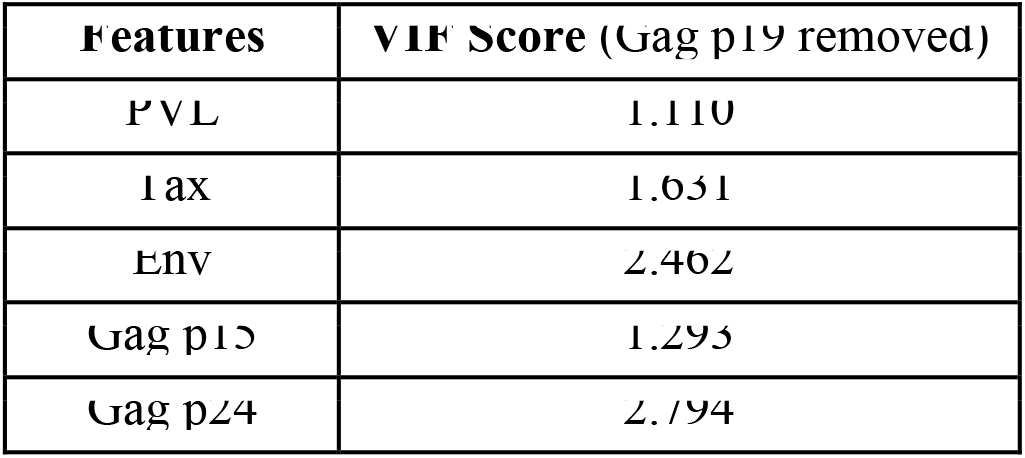
This table shows the reduction in VIF scores across all features after removing Gag p19. Recalculations of the VIF scores for the remaining features demonstrated a significant reduction in multicollinearity with all features displaying VIF-score < 3, suggesting that the remaining set of features PVL, Tax, Env, Gag p15 and p24 exhibits minimal multicollinearity and suitable for further analysis [O’Brien, R. M.].

**Supplementary Table S2:** Spearman’s rank correlation coefficients (r) illustrate the relationships between the feature values of the anomaly carriers (n=17) and their corresponding anomaly scores as determined by the Isolation forest anomaly detection model. * indicates a statistically significant correlation (p-value <0.05).

| Features | Spearman's rank correlation coefficient (r) among the feature values and the anomaly scores of the anomaly carrier data | p-value |
| --- | --- | --- |
| Gag p15 | -0.67 | 0.003* |
| Gag p24 | -0.55 | 0.022* |
| Env | -0.52 | 0.032* |
| Tax | -0.27 | 0.299 |
| PVL | 0.1 | 0.701 |

**Supplementary Table S3:** Overall Mean PRAUC of the classifiers used in our study.

| Classifier names | Overall Mean PRAUC<br>(One-vs-Rest multiclass classification) |
| --- | --- |
| Random Forest (RF) | 0.841 |
| XGBoost (XGB) | 0.819 |
| Extra Tree Classifier (ETC) | 0.839 |
| Support Vector Machine (SVM) | 0.811 |

**Supplementary Table S4:**
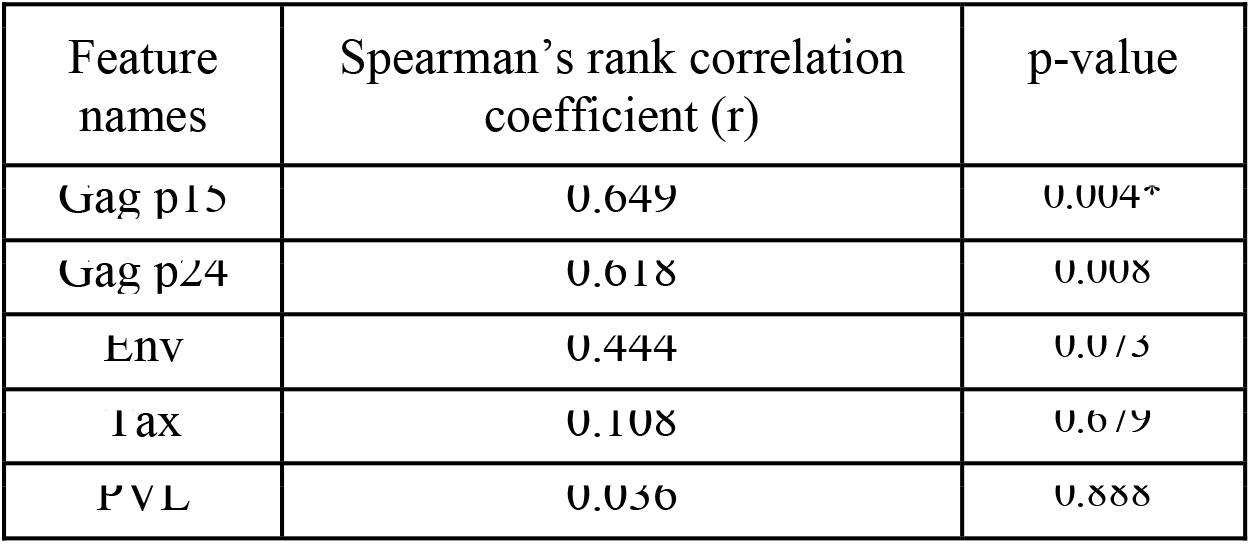
Spearman’s rank correlation coefficients (r) between the feature values of the anomaly carriers (n=17) and their predicted probability of belonging to HAM/TSP, as determined by the random forest classifier model. All the feature values except PVL showed a stronger positive correlation with the predicted probability of HAM/TSP. However, none of these were statistically significant except Gag p15. * indicates a statistically significant correlation (p-value <0.05).

**Supplementary Table S5:** This table represents the HTLV-1 Pro-viral load (PVL) and the antibody titer profile of the anomaly carrier samples with their respective anomaly scores obtained from the Isolation forest anomaly detection model. The sklearn implementation of the decision function of Isolation Forest yields anomaly scores, where negative scores indicate potential anomalies and positive scores indicate non-anomalies.

| Carrier ID | PVL (%) | Tax | Env | Gag p15 | Gag p24 | Anomaly Score |
| --- | --- | --- | --- | --- | --- | --- |
| indiv0027 | 35.1 | 46360.6 | 8036.57 | 133435.5 | 9186793 | -0.227 |
| indiv0035 | 46.3 | 6257.36 | 3380.86 | 1392 | 557465.7 | -0.107 |
| indiv0069 | 2.73 | 5475.765 | 6935.648 | 334938.483 | 10298208.9 | -0.206 |
| indiv0086 | 2.8 | 85900.225 | 4353.48 | 9175.32 | 7825533.16 | -0.135 |
| indiv0125 | 3.52 | 24760.56 | 9624 | 143221.032 | 10627154.2 | -0.143 |
| indiv0131 | 1.89 | 9997.35 | 10307.304 | 156990.407 | 8509020.98 | -0.110 |
| indiv0133 | 1.54 | 6710.417 | 2773.046 | 62421.852 | 7597349.42 | -0.051 |
| indiv0169 | 2.79 | 73261.68 | 10899.912 | 156270.114 | 5645401.27 | -0.160 |
| indiv0189 | 0.36 | 112722.516 | 2923.854 | 365.62 | 14694.135 | -0.113 |
| indiv0217 | 7.09 | 40165.452 | 8748.922 | 232614.22 | 5608868.82 | -0.136 |
| indiv0221 | 9.79 | 5629.212 | 14189.266 | 89869.177 | 5905812.24 | -0.133 |
| indiv0227 | 7.16 | 65582 | 14497.938 | 264498.926 | 7607445.6 | -0.215 |
| indiv0240 | 7.49 | 12044.324 | 9279.452 | 127028.479 | 8400371.16 | -0.087 |
| indiv0245 | 23.47 | 1863.456 | 3303.802 | 44529.681 | 1584100.75 | -0.051 |
| indiv0249 | 11.55 | 76016.766 | 3491.858 | 8507.275 | 665555.814 | -0.072 |
| indiv0254 | 5.07 | 28942.788 | 5466.446 | 335210.855 | 10180285.4 | -0.204 |
| indiv0265 | 19.76 | 29540.5446 | 7478.61089 | 65475.1918 | 4953871.65 | -0.107 |

**Supplementary Table S6:** Dunn’s post-hoc analysis followed by the Kruskal-Wallis test. This table demonstrates statistical differences between pairwise comparisons of the sample groups.

| Pairwise comparisons | PVL | Tax | Env | Gag p15 | Gag p19 | Gag p24 |
| --- | --- | --- | --- | --- | --- | --- |
| Carrier vs Anomaly carrier | Significant<br>P= 0.0197 | Significant<br>P= 0.0000 | Significant<br>P=0.0000 | Significant<br>P=0.0000 | Significant<br>P=0.0000 | Significant<br>P=0.0000 |
| HAM vs Anomaly carrier | Not significant<br>P= 1.0000 | Not significant<br>P= 1.0000 | Significant<br>P=0.0048 | Not significant<br>P= 0.3395 | Not significant<br>P= 0.6504 | Not significant<br>P= 0.8887 |
| ATL vs Anomaly carrier | Significant<br>P= 0.0190 | Significant<br>P=0.0001 | Significant<br>P=0.0000 | Significant<br>P=0.0000 | Significant<br>P=0.0446 | Significant<br>P=0.0137 |
| Carrier vs HAM | Significant<br>P= 0.0000 | Significant<br>P=0.0000 | Not significant<br>P= 0.3102 | Significant<br>P=0.0000 | Significant<br>P=0.0000 | Significant<br>P=0.0000 |
| Carrier vs ATL | Significant<br>P= 0.0000 | Not Significant<br>p= 1.0000 | Not significant<br>P= 1.0000 | Not significant<br>P= 1.0000 | Significant<br>P=0.0054 | Significant<br>P=0.0095 |
| HAM vs ATL | Significant<br>P= 0.0000 | Significant<br>P=0.0000 | Not significant<br>P= 0.1995 | Significant<br>P=0.0000 | Not significant<br>P= 0.6865 | Not significant<br>P= 0.1148 |

**Supplementary Table S7:**
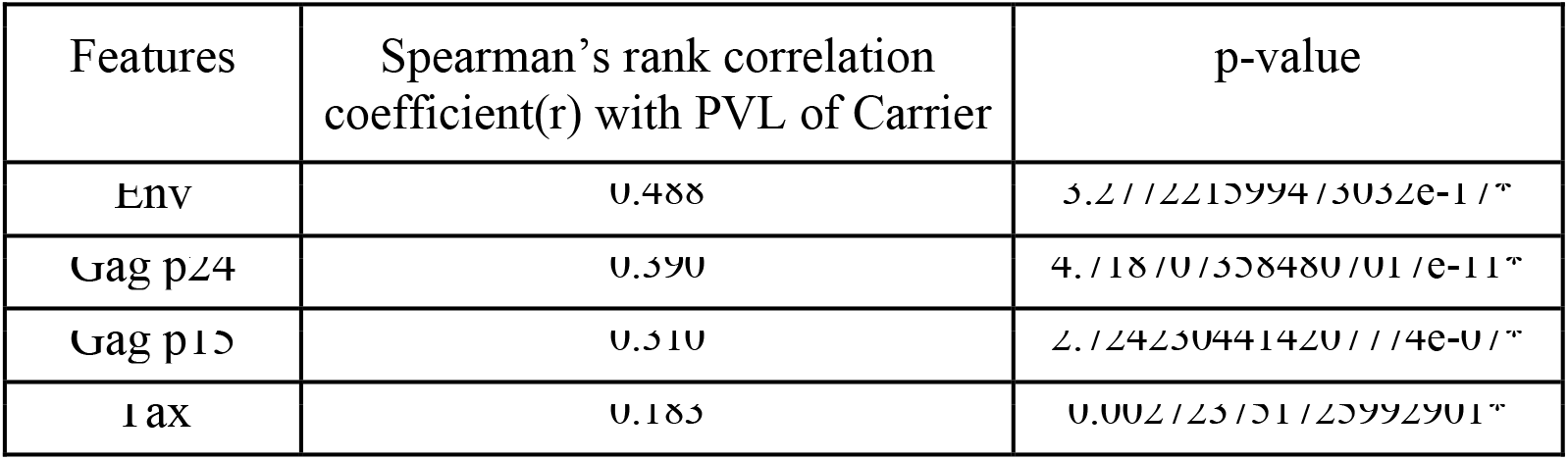
Spearman’s rank correlation analysis between PVL and other features among the Carrier population (n=264). * indicates a statistically significant correlation (p-value <0.05).

**Supplementary Table S8:**
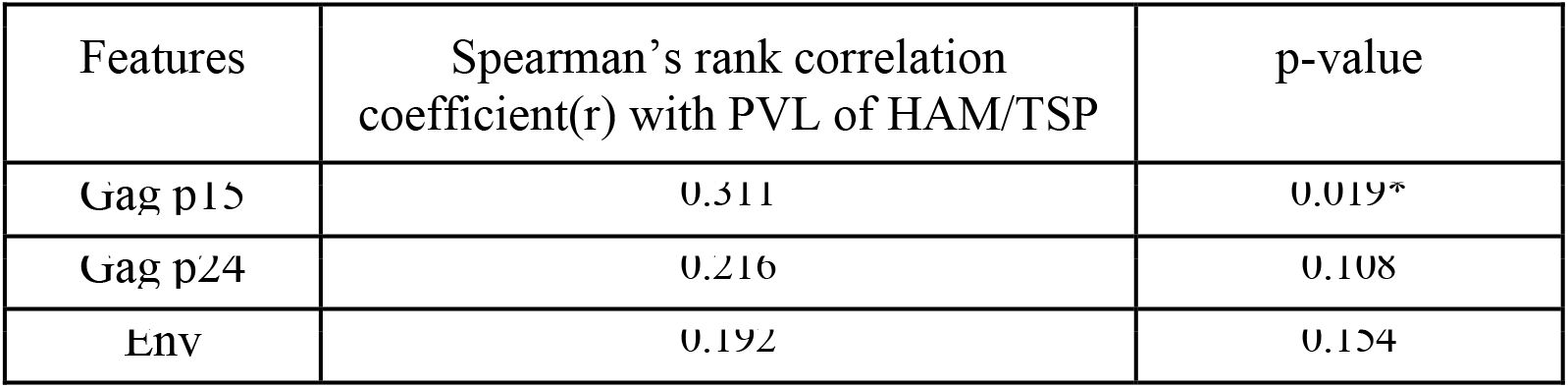
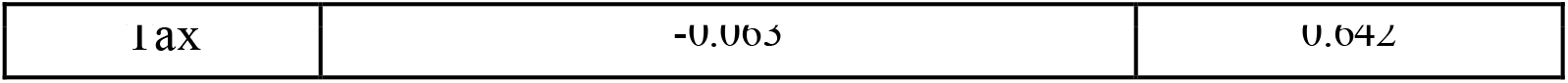
Spearman’s rank correlation analysis between PVL and other features among the HAM population (n=56). * indicates a statistically significant correlation (p-value <0.05).

**Supplementary Table S9:**
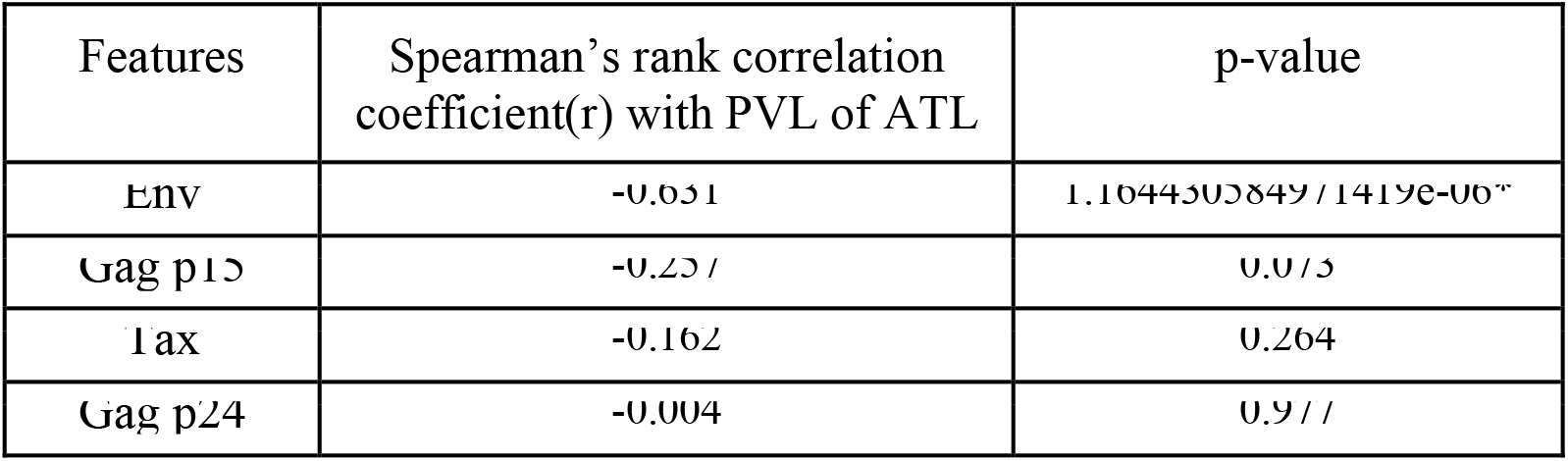
Spearman’s rank correlation analysis between PVL and other features among the ATL population (n=49). * indicates a statistically significant correlation (p-value <0.05).

### Supplementary Figures

**Supplementary Figure S1:**
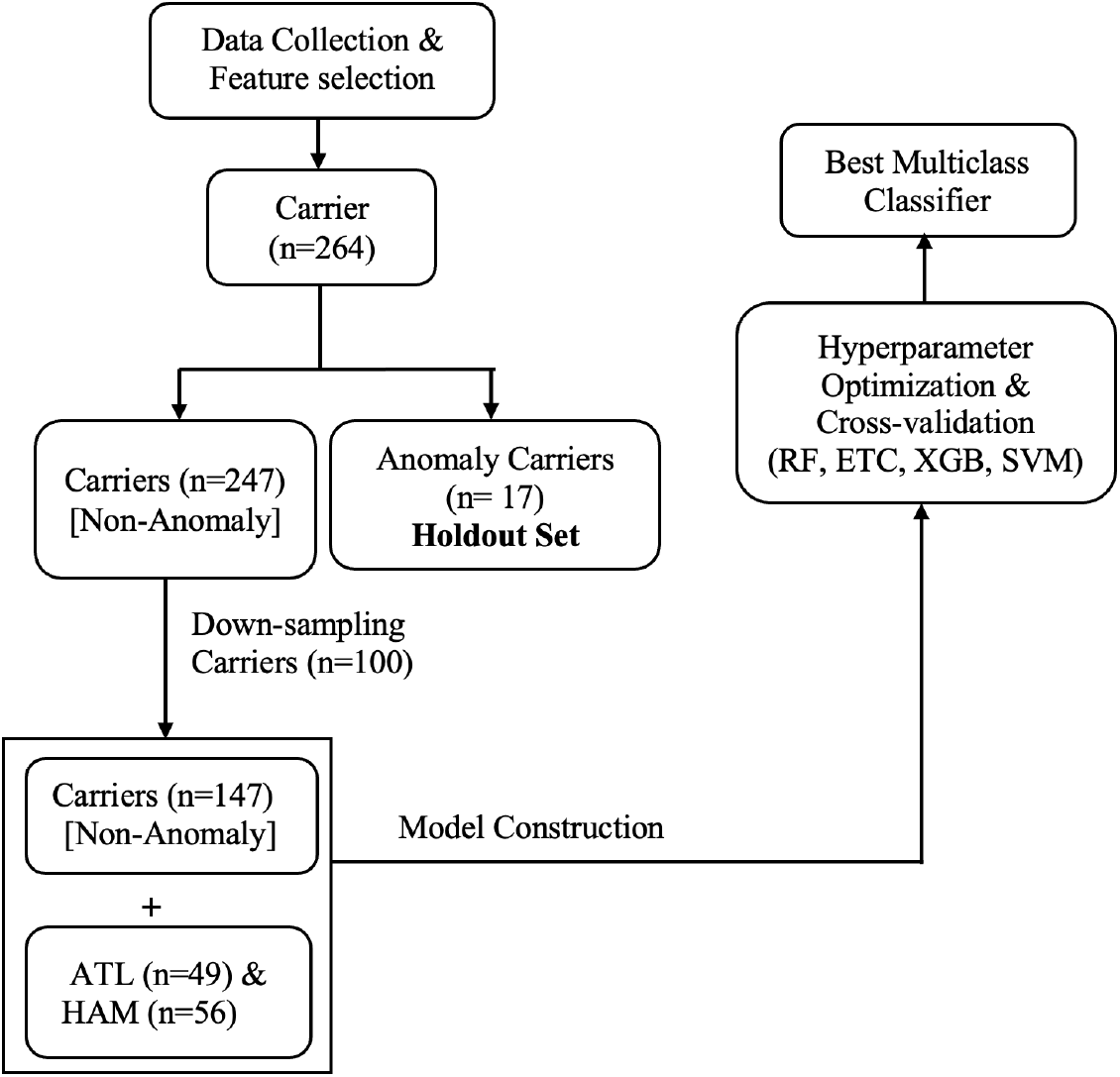
Workflow chart

**Supplementary Figure S2:**
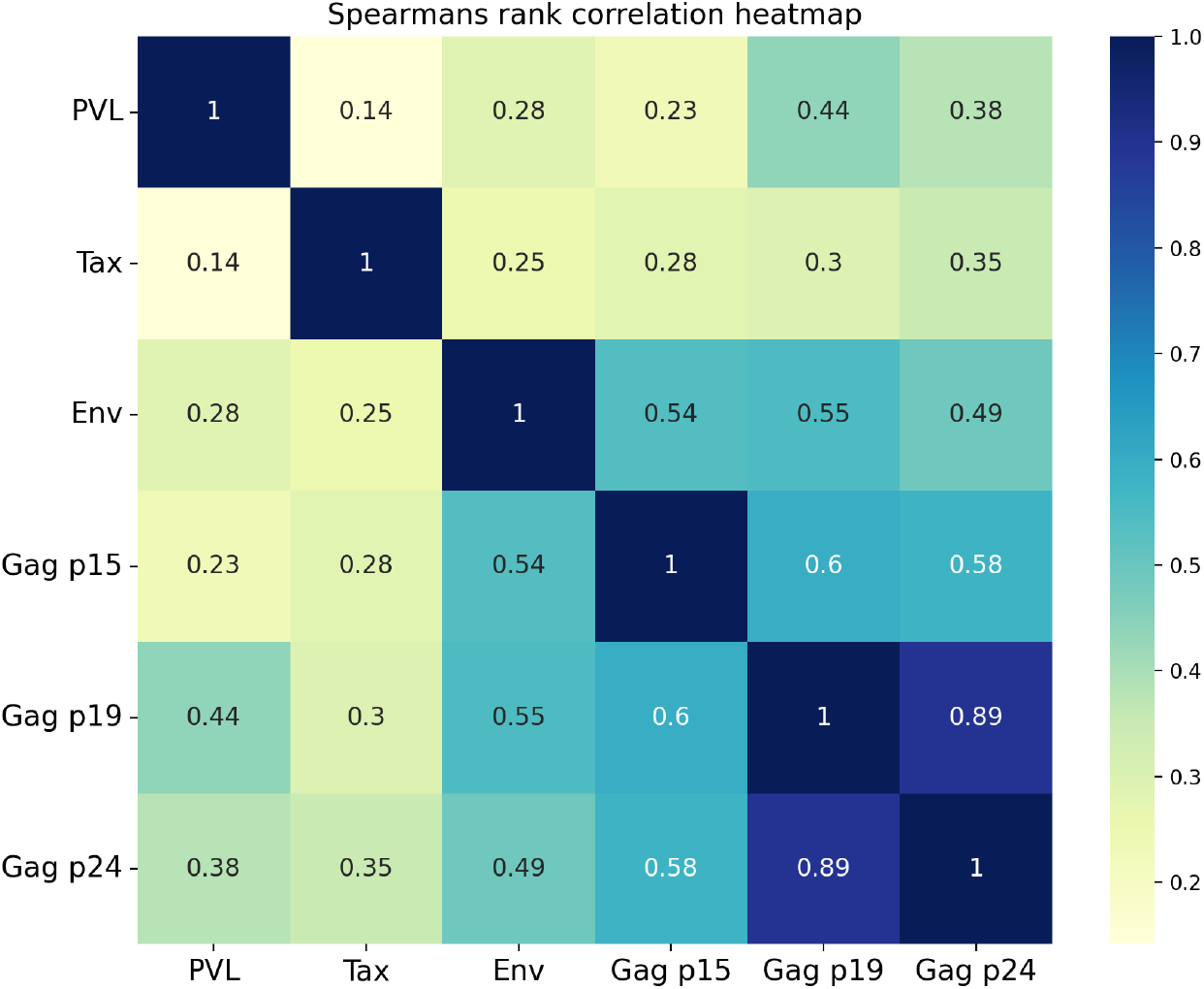
Spearman rank correlation heatmap among the feature values of all samples.

**Supplementary Figure S3:**
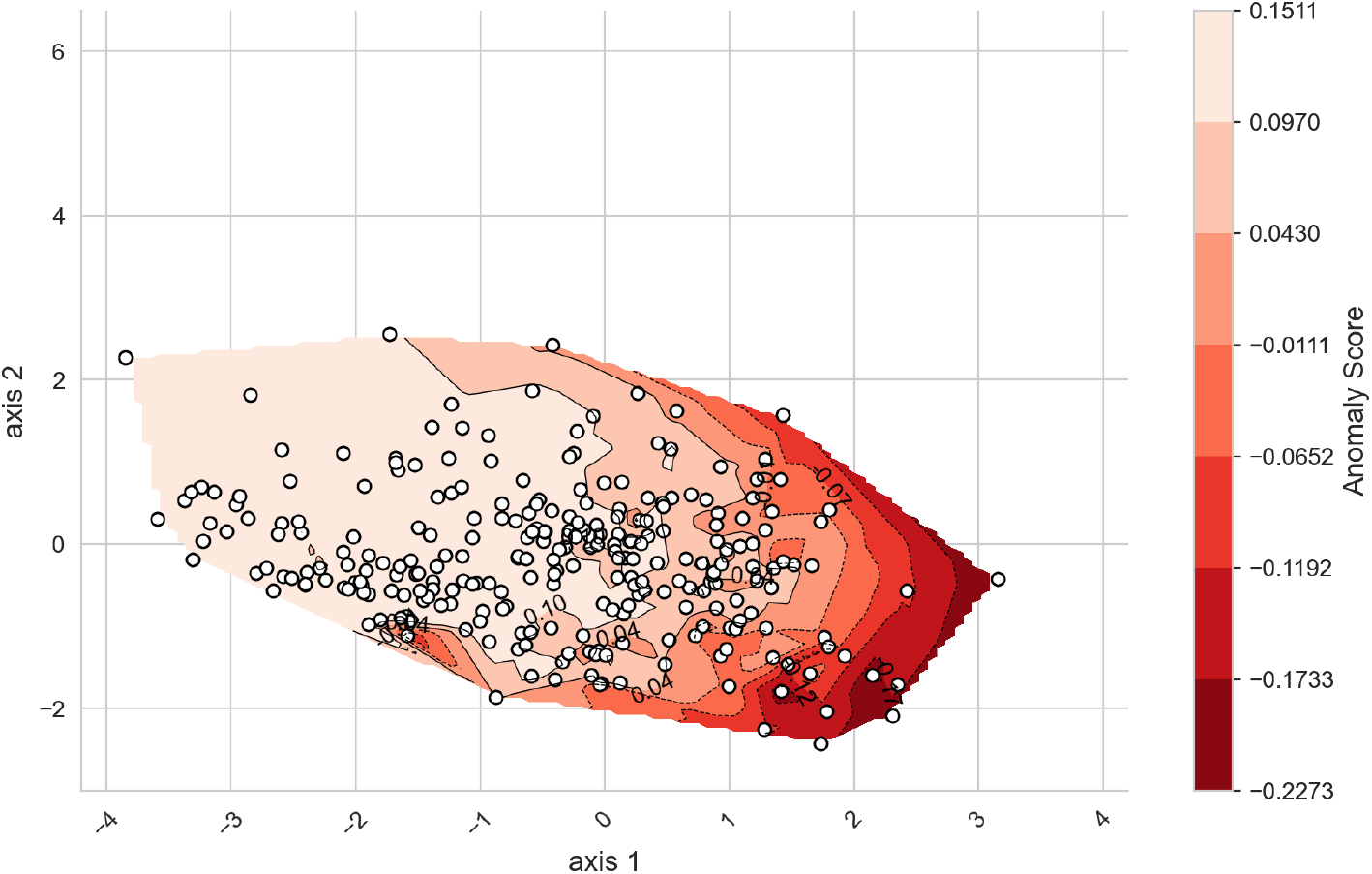
The contour plot provides a visual representation of the anomaly score distribution for the carrier-only data. Data points are higher in the lighter red areas, suggesting that most carriers have similar characteristics or behaviors, indicating they are less anomalous. In contrast, the points in regions with higher anomaly scores (darker shades) are more likely to be anomalies or outliers. The spread along these axes indicates how the data varies across these two dimensions. The color bar on the right provides a scale for the anomaly scores. Higher anomaly scores (darker shades) indicate a higher likelihood of being an anomaly.

**Supplementary Figure S4:**
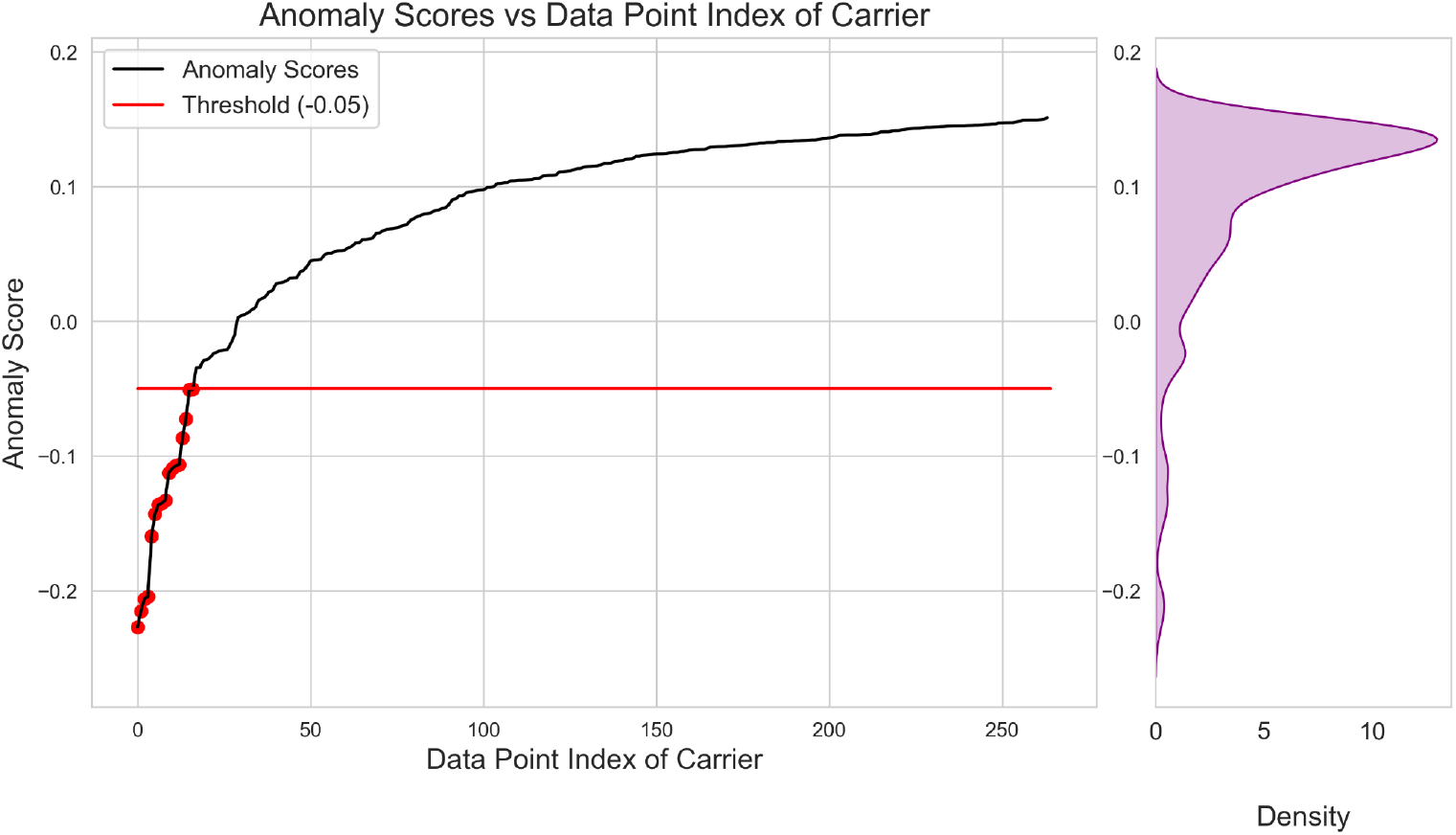
This figure shows the Isolation forest anomaly scores plotted against the carrier data point index (n=264) with a threshold of -0.05. The plot demonstrates that most data points have anomaly scores above the threshold (−0.05). Data points below the threshold were considered anomaly samples in our study and thus highlighted in red circles. The vertical figure on the right demonstrates the KDE distribution plot of the anomaly scores among the carrier population.

**Supplementary Figure S5:**
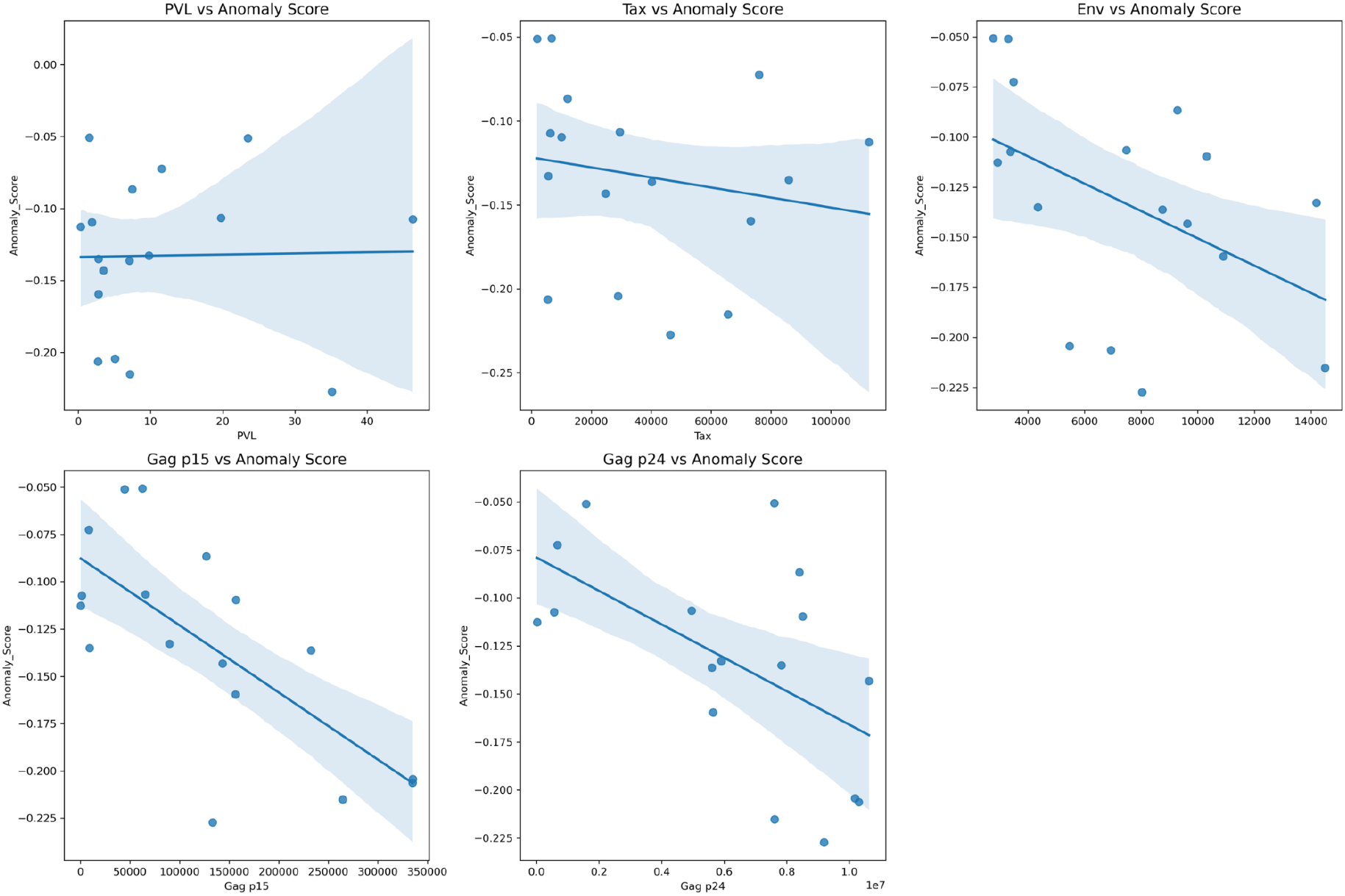
All features except PVL show a negative correlation with their respective feature values, indicating that as the feature value increases, the anomaly score tends to decrease (in sklearn implementation of Isolation forest, negative anomaly scores indicate anomalies). The strength of the negative correlation varies among the features, with Gag p15 showing the most substantial negative correlations compared to other features. The plots collectively suggest that higher values in the features Tax, Env, Gag p15, and Gag p24 are generally associated with lower anomaly scores. The shaded area indicates a 95% confidence interval.

**Supplementary Figure S6:**
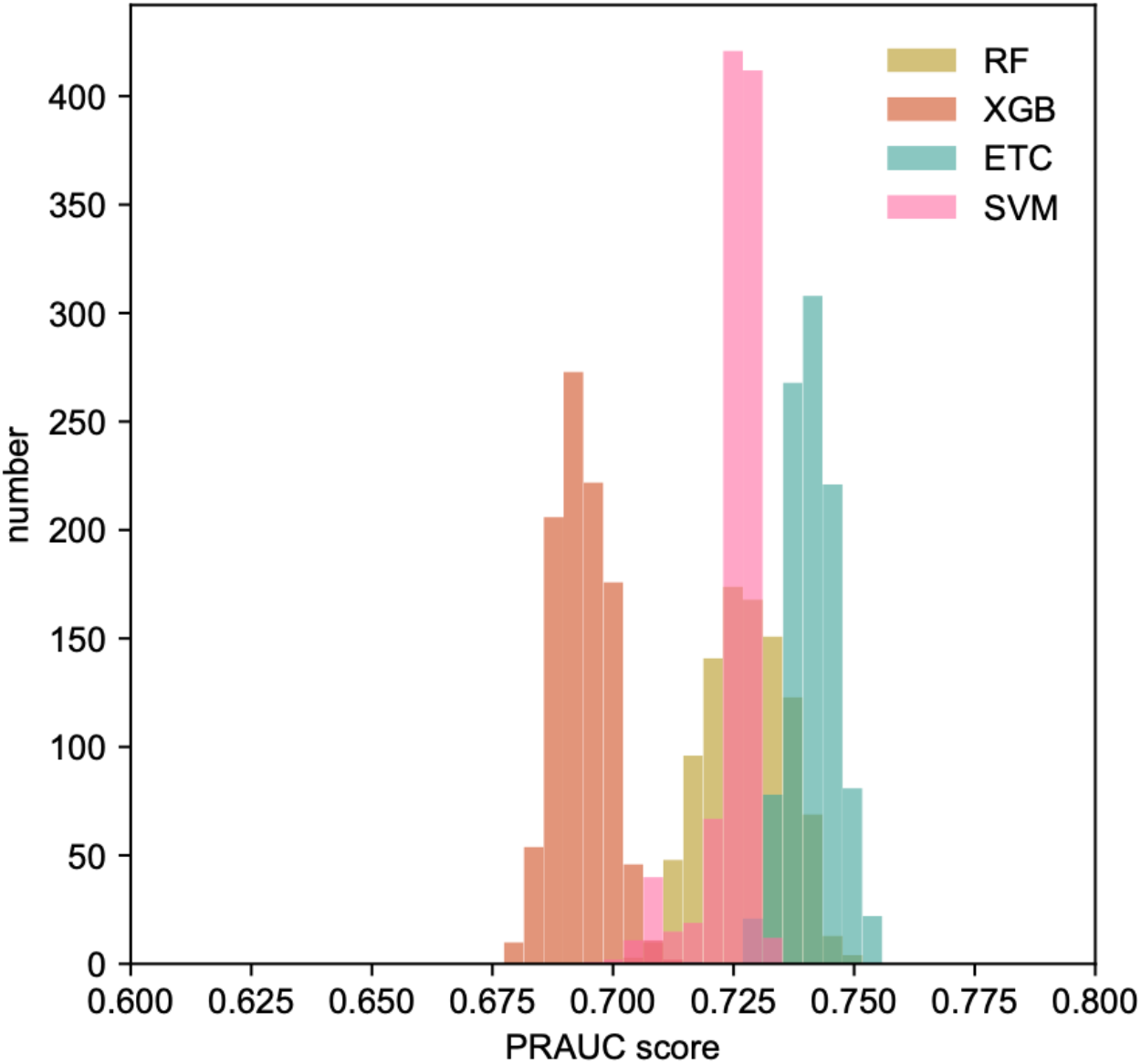
Histogram of performance for each classifier in terms of PRAUC. Each distribution was made from 1000 iterations of different random seeds (random_state parameter). In this study, 1000 patterns of random seeds are considered to evaluate the performance with a high degree of confidence. These results obtained from different random seeds then turn out to be reliable whenever the randomized manipulation during the learning process is different.

**Supplementary Figure S7:**
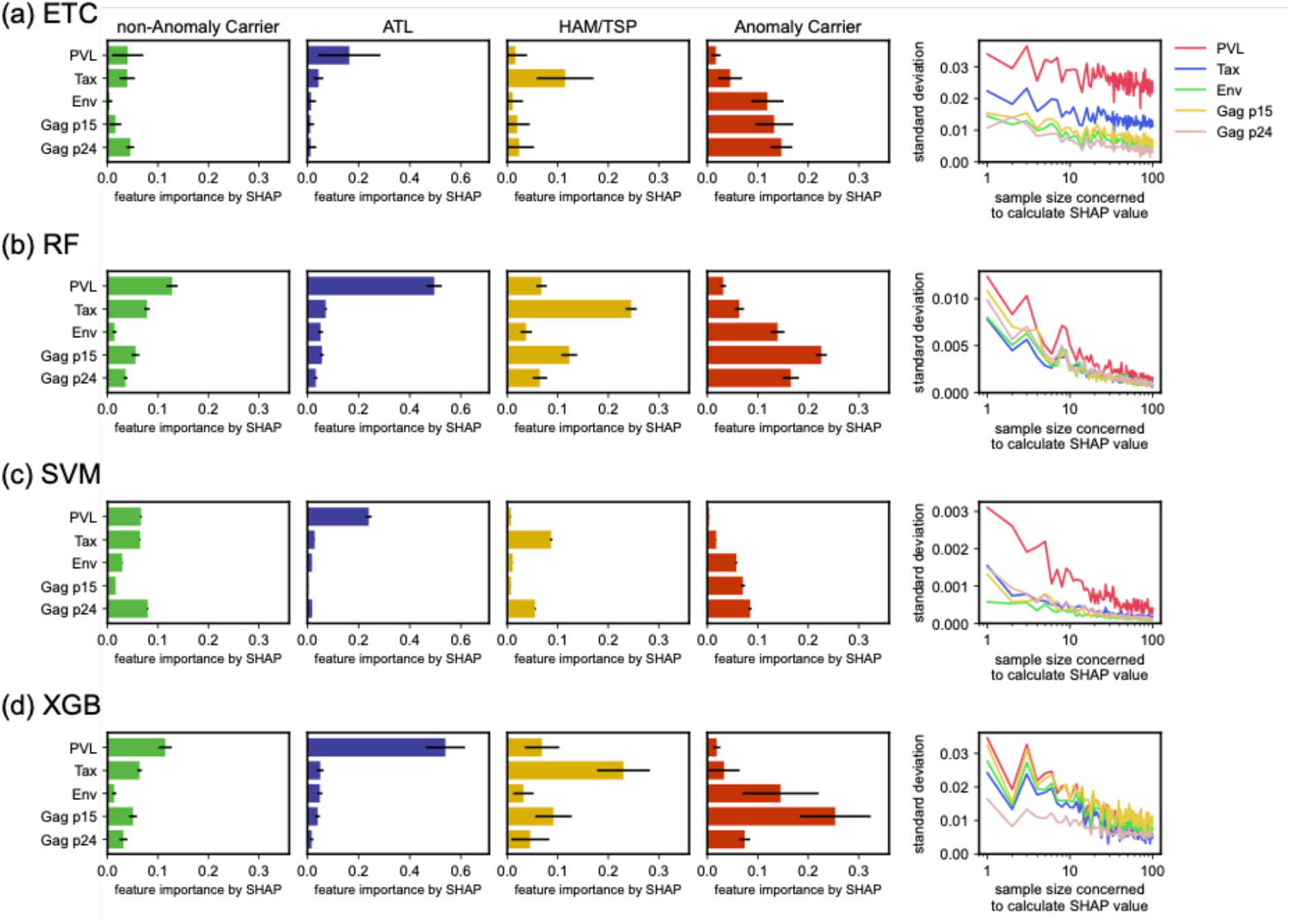
SHAP bar plot for each subgroup (non-anomaly carrier, ATL, HAM/TSP, and anomaly carrier) for each classifier (ETC, RF, SVM, and XGB) (left four panels). Each SHAP bar is a median value from 1000 iterations of different random seeds and the black line represents a standard deviation. The most right panel shows the standard deviation calculated from ten sets of different median SHAP values for each feature. In each set, a random seed is randomly picked from 1 to 100 and the median value is chosen (this is the x-axis). The figure showed that 100 iterations can converge to settle near zero; thus the SHAP bar plot was calculated using 1000 iterations (10 times higher). While the best-performing classifier was ETC [Supplementary Figure S6], the PRAUC scores of the four classifiers were comparable. Hence, we stuck to the idea that only consistent trends in SHAP values among all classifiers were considered reliable (SHAP values were consistent among all classifiers except for Gag p24).

